# Elicitation of potent SARS-CoV-2 neutralizing antibody responses through immunization using a versatile adenovirus-inspired multimerization platform

**DOI:** 10.1101/2021.09.13.460076

**Authors:** C Chevillard, A Amen, S Besson, D Hannani, I Bally, V Dettling, E Gout, CJ Moreau, M Buisson, S Gallet, D Fenel, E Vassal-Stermann, G Schoehn, P Poignard, MC Dagher, P Fender

**Affiliations:** CNRS, Univ. Grenoble Alpes, CEA, UMR5075, Institut de Biologie Structurale, 38042 Grenoble, France; Univ. Grenoble Alpes, CNRS, UMR 5525, VetAgro Sup, Grenoble INP, TIMC, 38000 Grenoble, France; CHU Grenoble Alpes, 38000 Grenoble, France; The Scripps Research Institute, La Jolla, CA 92037, USA

## Abstract

The severe acute respiratory syndrome coronavirus 2 (SARS-CoV-2) pandemic has shown that vaccine preparedness is critical to anticipate a fast response to emergent pathogens with high infectivity. To rapidly reach herd immunity, an affordable, easy to store and versatile vaccine platform is thus desirable. We previously designed a non-infectious adenovirus-inspired nanoparticle (ADDomer), and in the present work, we efficiently decorated this original vaccine platform with glycosylated receptor binding domain (RBD) of SARS-CoV-2. Cryo-Electron Microscopy structure revealed that up to 60 copies of this antigenic domain were bound on a single ADDomer particle with the symmetrical arrangements of a dodecahedron. Mouse immunization with the RBD decorated particles showed as early as the first immunization a significant anti-coronavirus humoral response, which was boosted after a second immunization. Neutralization assays with spike pseudo-typed-virus demonstrated the elicitation of strong neutralization titers. Remarkably, the existence of pre-existing immunity against adenoviral-derived particles enhanced the humoral response against SARS-CoV-2. This plug and play vaccine platform revisits the way of using adenovirus to combat emergent pathogens while potentially taking advantage of the adenovirus pre-immunity.

## Introduction

The severe acute respiratory syndrome coronavirus 2 (SARS-CoV-2), originated in Wuhan, China in 2019 causes severe pneumonia with 4 574 089 deaths as of September 8, 2021 (https://covid19.who.int/) and enormous socio-economic consequences^1^. SARS-CoV-2 is an enveloped positive strand RNA virus belonging to the beta-coronavirus genus from the coronaviridae family. On its membrane surface, it exposes the membrane heterotrimeric glycoprotein protein S, known as the Spike. The ectodomain of the S protein is divided into S1 and S2 domains. Both SARS-CoV-2 and SARS-CoV Spike proteins bind to a host Angiotensin-Converting Enzyme 2 (ACE2), which serves as an entry receptor. A subdomain of the S1 domain, named receptor binding domain (RBD)^2,3^, is the contact interface between the virus and the ACE2 receptors. Several neutralizing antibodies (NAbs)^4^ and decoy receptors^5^ have shown strong inhibition of cell infection by SARS-CoV-2 by competing with the RBD-ACE2 interaction.

The cryo-EM structure of the receptor/spike complex enabled to define precisely the key residues involved in the contacts and showed that their mutations altered binding affinity^2,3,6,7^. Due to its surface-exposed location and critical role in entry, the S protein is a main target of NAbs upon infection and therefore a good candidate for vaccine design. Thus, most vaccines currently licensed for humans use this protein as an antigen. Vaccine platforms are essentially based on the use of recombinant adenoviral vectors or mRNA. Adenovirus is an attractive non-replicative vector because of its ease of production and storage and was used in several of the currently authorized COVID vaccines^8^. However, pre-existing immunity in the human population is a potential concern to the effectiveness of such delivery vehicles resulting in the use of a non-human derived adenoviral vector or alternating use of several human serotypes such as HAdV-5 and HAdV-26^8^.

Beside the whole S protein, numerous studies have shown that the RBD comprises multiple distinct antigenic sites and is a prime target for neutralizing antibodies in COVID-19 convalescent patients^9–12^. Moreover, these NAbs have been shown to confer protection against SARS-CoV-2 challenge in animal models of COVID-19, as well as to prevent COVID-19 in human, thus confirming the interest of using RBD as a vaccine immunogen^13,14,15^.

We have previously reported that a non-infectious particle derived from human Adenovirus of type 3 consisting of 60 identical penton base monomers could be modified to display epitopes of interest including those from emerging viruses as its surface^16–18^. In this vaccine platform named ADDomer, exposed loops of the penton base protein were engineered to allow insertion of foreign peptides such as a linear neutralizing epitope from Chikungunya virus. However, insertion of structurally complex antigens was not permitted by this first platform. To overcome this limitation while keeping the immunological advantages of ADDomer, we describe here a novel adenovirus-inspired vaccine platform offering a unique highly versatile display of large and structurally complex antigens with post-translational modifications. To allow the versatile insertion of large antigens, the Spy Tag/Spy Catcher system^19–22^ was combined to the ADDomer technology. In this system derived from *Streptococcus pyogenes* an Aspartate residue from the 13 amino-acid SpyTag peptide (ST) can spontaneously create a covalent bond with a Lysine residue encompassed in the complementary SpyCatcher module (SC). We genetically inserted the sequence coding the ST peptide into the ‘variable loop’ of the ADDomer that was previously used for small antigen insertion. On the other hand, the RBD antigen from SARS-CoV-2 virus was fused to SC, in order to make a novel and original ADDomer-based COVID-19 vaccine. Here, we report that the newly designed platform can successfully be decorated with SARS-CoV-2 RBD and is highly immunogenic in mice, eliciting high neutralizing Ab titers.

## Results

### Internal insertion of ST in ADDomer results in efficient SC cross-linking

ADDomer is a non-infectious 30 nm nanoparticle, formed from 12 bricks of the homo-pentameric penton base from the human adenovirus type 3. The ST sequence was inserted in the ADDomer gene in a region coding for the exposed flexible loop called the variable loop (Figure 1a). Due to the spontaneous homo-oligomerization of twelve pentameric penton base, sixty copies of ST are exposed on the surface of the ADDomer-ST (ADD-ST) particle. To assess whether the newly inserted ST sequence was accessible and functional on ADD-ST, incubation with different ratio of SC was performed. SDS-PAGE profile clearly showed that the incremental SC/ADD-ST ratio is correlated with the apparition of a band of higher molecular weight and the decrease of the intensity of bands related to unlinked moieties. Importantly, there was no remaining signal for undecorated ADD-ST after SC addition at the highest ratio thus reflecting the full decoration of the nanoparticle (Figure 1b). This result suggests an efficient and concentration-dependent formation of the ADD-ST/SC complex. Mass spectroscopy analysis confirmed that the ADD-ST doublet shifted by 13,3kDa (Figure 1c), which corresponds to the molecular weight (MW) of SC. The two peaks observed in mass spectroscopy is due to a second initiation codon in ADD-ST (starting at Methionine 25) and SC can bind to each form indifferently as shown by a similar shift of their MW corresponding to the addition of SC. Negative staining electron microscopy imaging performed on ADD-ST alone and ADD-ST fully decorated by SC showed that the integrity of the particle was not affected by the presence of SC and the grainy appearance of ADD-ST/SC likely reflects the presence of SC at the particle surface (Figure 1d). Altogether, these data show that SC cross-linking to ADD-ST is operational and can result in saturated decoration of the nanoparticle.

**Figure 1:**
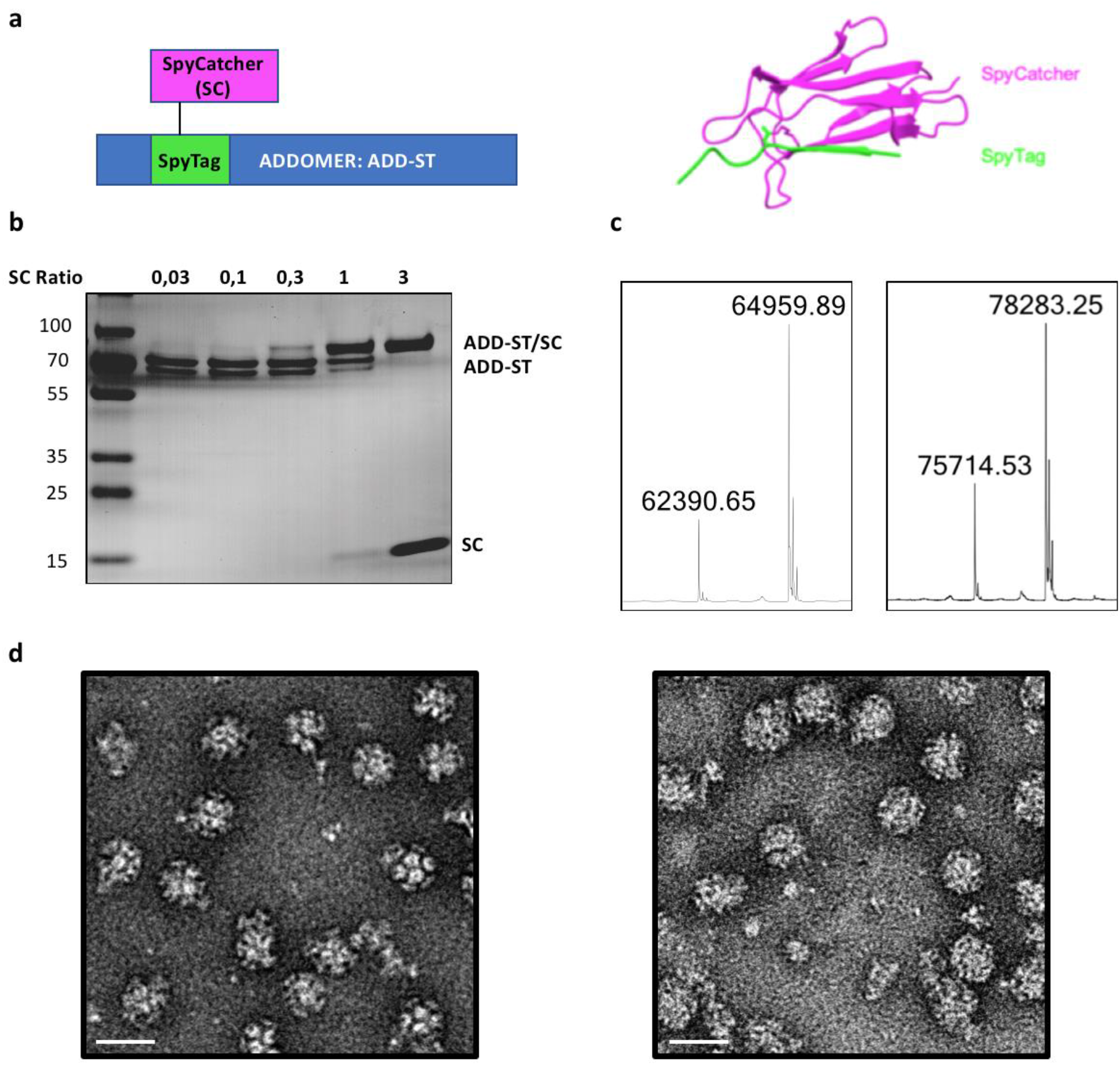
Application of the SpyTag/SpyCatcher cross-linking method to the ADDomer technology. (a) Diagram showing the internal insertion of SpyTag in the internal loop of an ADDomer monomer. Spytag (in green) can make an isopeptidic bond with SpyCatcher (in purple). Structure of a SpyTag covalently linked to a SpyCatcher is shown on the right (PDB code: 4MLI) (b) SDS-PAGE profile of ADD-ST interaction with incremental ratio of SC showing the apparition of a higher MW covalent adduct (ADD-ST/SC). (c) Electrospray ionization graphs of ADD-ST (left) and ADD-ST/SC (right) showing the shift of the peaks by 13.3 kDa. The doublet is due to an alternative start of translation in ADD-ST and display the same mass shift of 13,3kDa. (d) Negative staining electron micrographs of ADD-ST (left) and ADD-ST/SC (right, bar: 30nm).

### Cryo-EM analysis enables to visualize SC decoration on ADD-ST

In order to visualize the SC arrangement around the ADDomer particle, a structural study by cryo-electron microscopy was used. Fully decorated ADD-ST/SC particles were imaged on a Glacios electron microscope (Figure 2a). Image analysis was performed and the obtained 3D structure was compared to the structure of an undecorated ADDomer particle (EMD-0198). An extra-density corresponding to SC bound to the ADDomer ‘variable loop’ was clearly visible in ADD-ST/SC both in the density slice of the 3D map (Figure 2b) and in the isosurface representation of the structure (Figure 2c, purple density). The resolution of the ADDomer particle is around 2.8 Å, allowing to clearly see the aminopeptidic chain. However, the ST/SC part is not rigidly attached to the ADDomer, which explain why the corresponding density is fuzzy, and the structure not better defined. The ADDomers have 2, 3 and 5 axis of symmetry, (the latter two being shown in Figure 2c), which is a characteristic of a dodecahedron meaning that SC are distributed accordingly. Interestingly, along the 3-fold axis SC are ∼ 4.7 nm apart which is close to the Receptor Binding Domain (RBD) in the trimeric spike protein of SARS-CoV-2 (∼ 4.1 nm), suggesting that the decorated particles would closely mimic the natural trimeric arrangement of RBDs (Figure 2d). Both the multivalency of SC displayed by the particle and the arrangements they have around the different symmetry axis could be an asset for vaccination purpose if an antigen is fused to this module.

**Figure 2:**
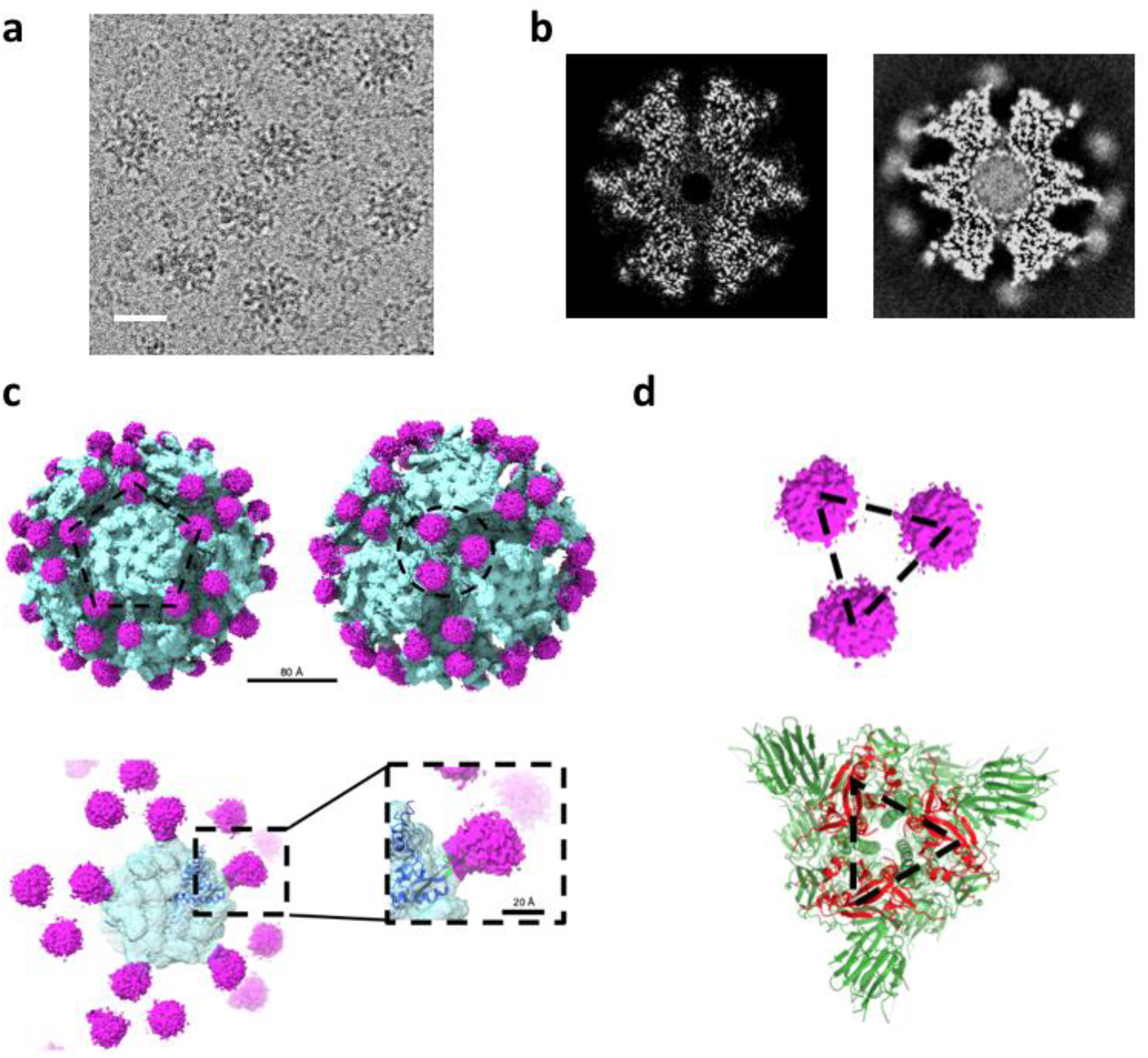
Cryo-EM reconstruction of ADD-ST decorated with SC. (a) Representative 2D picture of the particles frozen on ice (Bar:30nm). (b) Section through the density of the 3D reconstruction without or with SC (left and right, respectively). (c) Isosurface representation of the ADDomer-ST/SC 3D structure showing the extra density of SC in purple onto the non-decorated ADD-ST scaffold in light blue. The particle is represented along either the 5 or 3 fold-axis (left and right, upper panel; bars: 80 and 20Å, respectively). Focus on a pentameric complex of ADD-ST/SC with a close up view onto a single SC/ST interaction (dashed line boxes in the lower panel). The atomic resolution structure (dark blue) of the HAd3 penton base (PDB 4AQQ) has been fitted into the ADDomer EM density (light blue) (d) Organization of SC along the 3-fold axis of the particle (upper panel) and comparison with SARS-CoV-2 RBD architecture (in red) within the Spike protein structure (in green, PDB 6VXX) at the same scale.

### Secreted RBD fused to SC is glycosylated and can be displayed at different ratios on the ADD-ST particle

To take advantage of the spontaneous and covalent linking of SC to ADD-ST, we reasoned that SC could be used as a versatile carrier to easily and efficiently fuse epitopes, like in this SARS-CoV-2 vaccine development (Figure 3a), and in broader perspectives to diverse soluble proteins of interest. In the present study, the SARS-CoV-2 RBD was fused to the N-terminus of the SC (RBD-SC) and a melittin signal peptide was added to allow protein post-translational modifications (PTM) such as glycosylations and secretion from insect cells. To assess whether the secreted and purified RBD-SC was glycosylated, it was treated with N-glycosydase and the result showed a shift in the migration of the related band in SDS-PAGE gel, indicating that RBD-SC was indeed glycosylated (Figure 3b). As previously performed with unfused SC, incremental amounts of RBD-SC were added to ADD-ST in order to decorate the particle with different ratio of cargos (from 30 to 60 copies per particle). The SDS-PAGE gel profile (Figure 3c) showed the correlated apparition of a band at higher molecular weight with increased concentrations of RBD-SC. This result reflects the covalent linking of RBD-SC to the ADD-ST monomers whereas the band corresponding to non-decorated ADD-ST monomer progressively faded away. This result showed that the ADDomer platform can be decorated with different numbers of antigen copies exposed on its surface. Altogether, these experiments demonstrated that SC fusion to large antigens (∼40kDa for RBD-SC) with post-translational modifications enable their covalent linking to the ADDomer platform and that the ratio of decoration can be adjusted according to desired applications.

**Figure 3:**
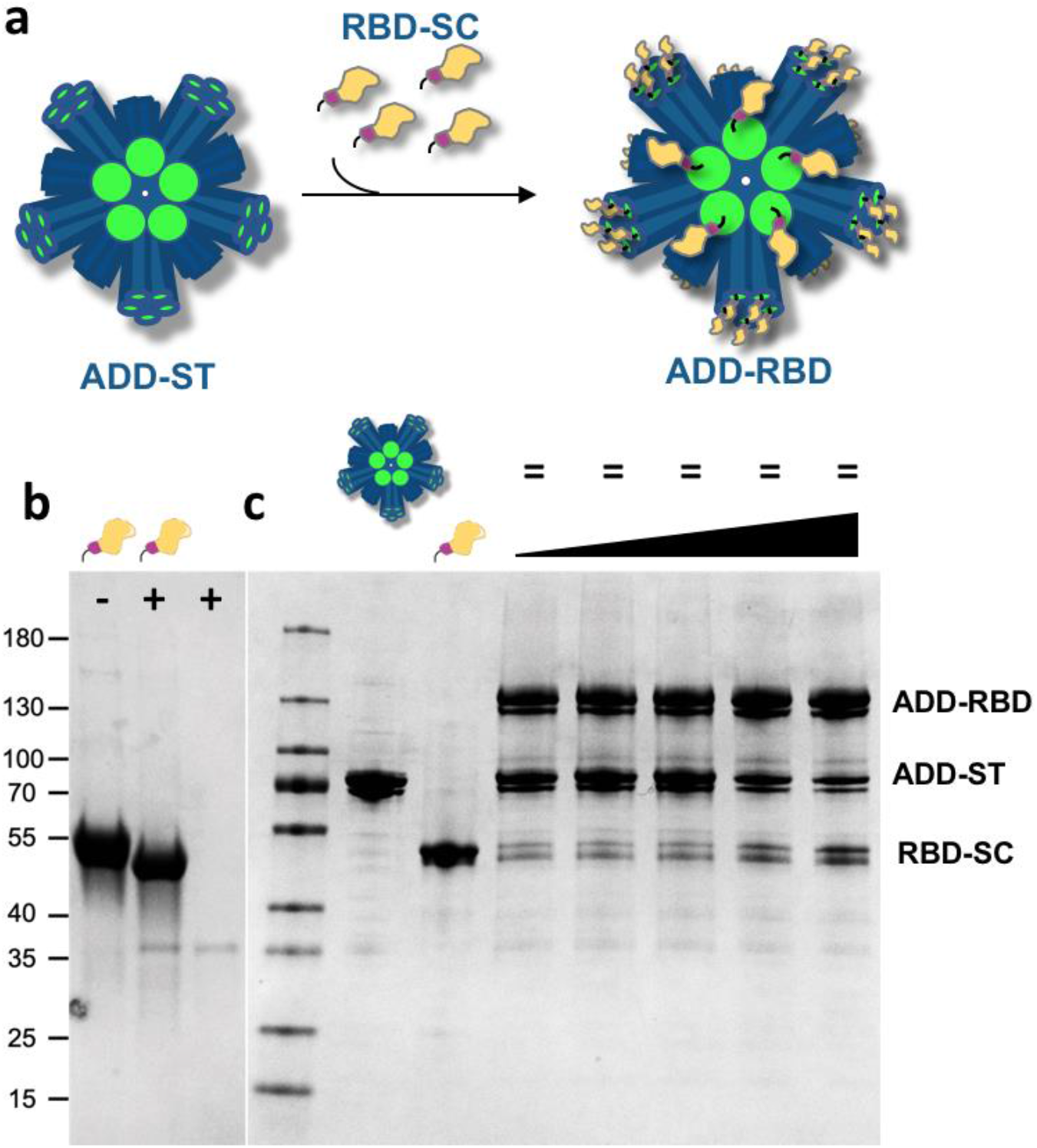
Fusion of SARS-CoV-2 RBD to SC enables its surface presentation on ADDomer particles. (a) Schematic representation of the spontaneous RBD-SC binding to ADD-ST to give ADD-RBD. (b) SDS-PAGE gel with RBD-SC before (-) and after (+) treatment with N-glycosidase and in absence of SC-RBD. The decrease of the molecular weight of SC-RBD after treatment indicates that it is glycosylated. (c) SDS-PAGE gel showing from left to right, the marker, bands of ADD-ST alone, SC-RBD alone and a fix amount of ADD-ST incubated with increasing amount of RBD-SC. The additional upper bands reflect the covalent adduct between ADD-ST and RBD-SC and thus the apparition of ADD-RBD. As expected, the increasing intensities of the ADD-RBD bands correlate with a decrease of non-decorated ADD-ST monomer.

### ADDomers decorated of glycosylated RBD bind to ACE2

RBD is the subdomain of the SARS-CoV-2 spike protein that binds to the human ACE2 receptor. To assess the function of RBDs linked to the ADDomer particles, three different approaches were used. First, binding experiments at the molecular scale were performed using surface plasmon resonance (SPR) on immobilized dimeric human ACE2 fused to Fc domain of human IgG^23^. Sensorgrams of ADD-RBD binding to immobilized human ACE2 showed a clear concentration-dependent binding of the RBD-decorated particles on ACE2 and a stable interaction with no visible dissociation at the end of injections (Figure 4a). This quasi-irreversible interaction agreed with a sub-nanomolar apparent K_D_ (3.09 × 10^−10^ M). In a second experiment at the cell scale, the direct binding of ADD-RBD onto ACE2-expressing HeLa cells was visualized by immunofluorescence using anti-ADDomer antibodies. The green signal seen at the periphery of HeLa-ACE2 cells in presence of ADD-RBD contrasted with the absence of signal observed with the same cells but in presence of undecorated ADD-ST particles. This result demonstrated that RBD at the surface of the particles induced the interaction with ACE2 present at the surface of the HeLa cells (Figure 4b). Finally, a competition experiment between a pseudo-typed virus harboring the SARS-CoV-2 spike and either RBD-SC or ADD-RBD was performed. Negative control was made with undecorated ADD-ST alone. As expected, both RBD-SC alone and ADD-RBD were able to compete with the pseudo-typed virus with a slightly higher efficiency for the RBD-decorated particle (Fig. 4c). Altogether, these experiments showed that RBD-SC is properly folded and can bind the ACE2 receptor at the molecular and cellular scales when applied alone (RBD-SC) or linked to the ADDomer particles (ADD-RBD).

**Figure 4:**
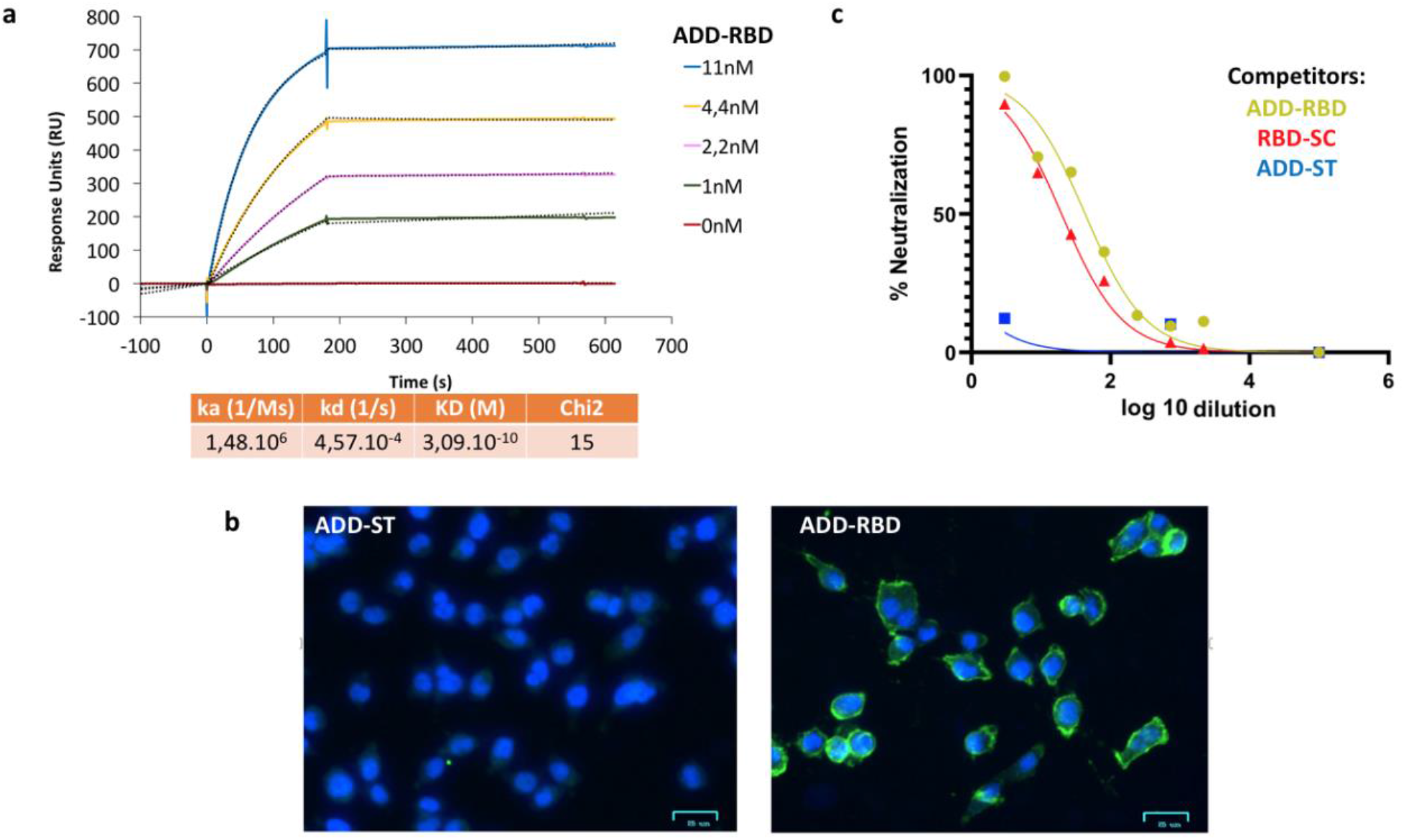
Functional characterization of the ADDomer particles decorated with SARS-CoV-2 RBD. (a) Surface plasmon resonance sensorgrams obtained by injection of different amounts of ADD-RBD to the immobilized ectodomain of the human ACE2 receptor fused to IgG Fc constant domain^23^ and kinetic analysis. (b) Immunofluorescence microscopy images of ACE-2 HeLa cell at 4°C with double staining with Hoechst (blue) and anti-ADDomer antibody and an Alexa488-conjugated secondary antibody (green) in presence of non-decorated ADD-ST (left) or ADD-RBD (right) particles. (c) Competition of pseudotyped SARS-CoV-2 virus encoding luciferase with either ADD-ST (blue), RBD-SC (red) and ADD-RBD (gold) at different dilutions.

### RBD-decorated ADDomer elicits rapid anti-CoV2 antibody responses in mice, enhanced by adenovirus pre-immunity

Multivalent exposition of RBD antigens at the surface of the particle is likely to result in a better activation of the humoral system than RBD alone. However, one cannot exclude that ADDomer itself could also play an indirect role in the anti-SARS-CoV-2 response, especially in a population with preexisting adenovirus immunity. To address these two points, four groups of ten mice were designed (Figure 5a). The two first control groups were injected with RBD-SC alone in the group 1 or with the same amount of unlinked RBD-SC plus naked-ADDomer (*i*.*e*. not displaying the antigen) in the group 2. The two other groups (groups 3 and 4) were vaccinated with ADD-RBD (*i*.*e*. RBD displayed at the particle surface) but the group 4 was pre-immunized with naked-ADDomer two weeks before the injection of ADD-RBD, to investigate the role of adenovirus pre-immunity (Figure 5b). The presence of anti-adenovirus antibodies in group 4 was checked one day before the first immunization with ADD-RBD by ELISA using anti-ADDomer antibodies. The results (Supplementary Figure 1) show that all mice of this group were preimmunized against naked-ADDomer. Then, the immune response against RBD was monitored by ELISA at day 13 (post-1^st^ immunization) and day 27 and 41 (post-2^nd^ immunization). The RBD-decorated particles induced a significant anti-SARS-CoV-2 response after the first immunization (day 13, figure 5c), whereas no response was detectable in controls of group 1 (RBD-SC alone) or group 2 (ADDomer with not displayed SC-RBD). Interestingly, adenovirus pre-immunity enhanced significantly the anti-RBD response.

**Figure 5 :**
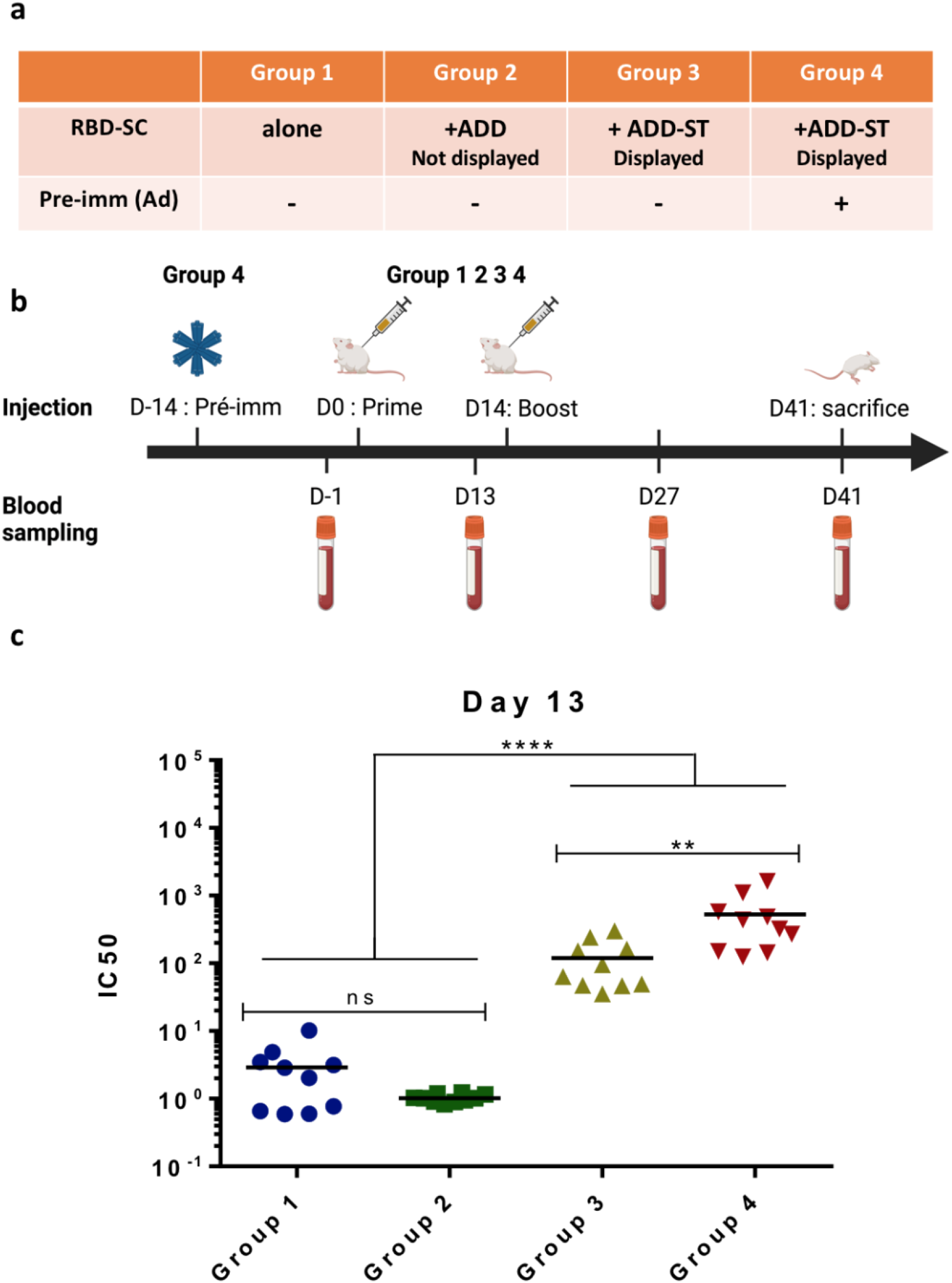
Immunization studies on mice inoculated with RBD-decorated ADDomer particles. (a) Different groups were constituted to address the respective role of RBD-SC alone (group 1), RBD-SC in presence of naked-ADDomer (group 2) and RBD-SC displayed by ADDomer (ADD-RBD) either in adenovirus naïve or adenovirus pre-immunized mice (group 3 and 4 respectively). (b) Immunization schedule and blood sampling for all groups. (c) IC50 of anti RBD response 13 days after the first immunization (n=10). Lines are mean values. Statistical analysis was performed by Prism.

### Immunization boost increases both the amplitude and the duration of the response induced by ADD-RBD particles, independently of the adenovirus immunological status

Two weeks after the second immunization (day 27), the anti-RBD response was clearly boosted for groups 3 and 4 that were injected with ADD-RBD, whereas the control groups 1 and 2 showed more heterogeneous responses (Figure 6a). This observation definitely confirms that our vaccine platform displaying multivalent RBD antigens yields a rapid and efficient humoral response. The ADDomer technology applied to RBD antigen also demonstrated a long lasting immunogenic effect with anti-SARS-CoV-2 RBD antibodies remaining at the highest level at day 41 (Figure 6b). The difference between mice pre-immunized by adenovirus (group 4) or not (group 3) was less pronounced than after the first immunization, both groups reaching high and comparable levels. Altogether, these results showed that the second immunization enabled a high and long-lasting antibody response against the SARS-CoV-2 RBD using the ADDomer platform and that adenovirus pre-immunity is beneficial in particular to the initial responses (Figure 6c).

**Figure 6:**
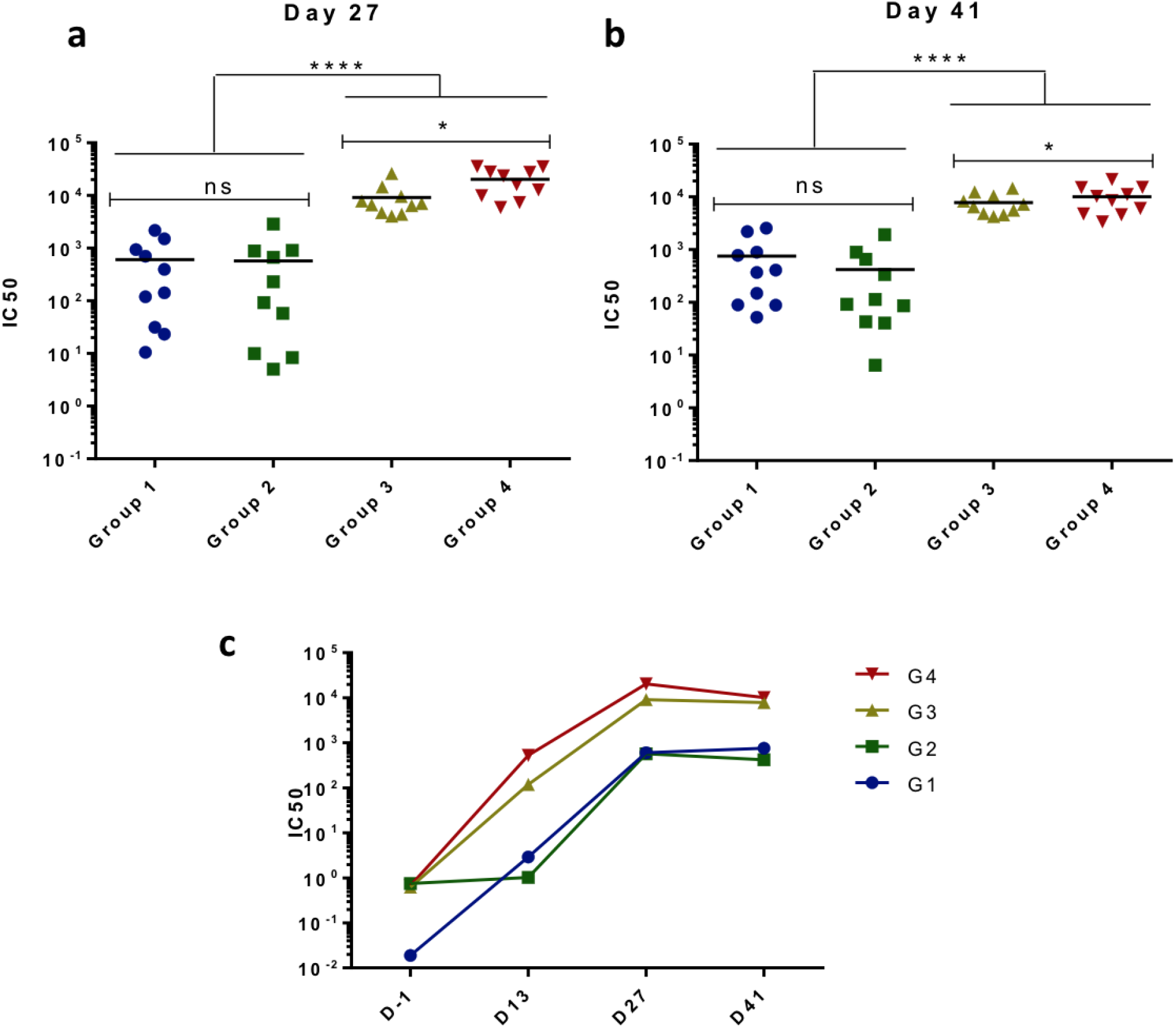
Anti-RBD antibody in all groups of mice 2 and 4 weeks after the booster immunization. (a) IC50 of anti-RBD for each individual mice from all groups, 2 weeks after the booster injection. (b) Same data, 4 weeks after the booster injection. Lines are mean values. Statistical analysis was performed by Prism. (c) Means of the anti-RBD response from all mice according to their groups and the time after the first immunization performed at Day 0.

### RBD-decorated ADDomer immunization elicits strong neutralizing antibody responses

Since the RBD is critical for binding to the SARS-CoV-2 receptor ACE2, antibodies elicited through immunization with the RBD (RBD-SC or ADDomer-RBD) can hinder the interaction between the Spike proteins and ACE2, blocking viral entry. The ability to neutralize the virus depends on the affinity of the antibodies for the RBD domain, as well as on the epitope recognized. To assess the potency and efficacy of the antibodies induced in the four immunized groups of mice, the neutralization potency of sera (day 41) from vaccinated mice were assessed using SARS-CoV-2 spike (Wuhan strain) pseudo-typed virus and ACE2-expressing HeLa cells. Only a partial and heterogeneous neutralization was obtained from groups 1 and 2 sera in which RBD-SC was not displayed on ADDomer even after two immunizations (Figure 7a, upper panels). Of note, sera from mice immunized with RBD-decorated ADDomer (groups 3 and 4) showed a better neutralization titer even after a single injection (Supplementary Figure 2). After the second immunization, strong virus neutralization was observed in all mice from groups 3 and 4 (Figure7a, lower panels). Neutralization titer calculation (ND_50_) clearly showed that mice vaccinated with RBD-decorated ADDomer had significantly higher neutralizing antibody titers than mice injected with the same amount of antigen but not displayed on the platform (Figure 7b). Moreover, in accordance with the ELISA results (Figure 5c), the adenovirus pre-existing immunity (group 4) slightly increased the neutralization titer compared to the naïve group (group 3), which emphasizes that this pre-existing adenovirus immunity appears beneficial for our novel vaccine platform (Figure 7b).

**Figure 7:**
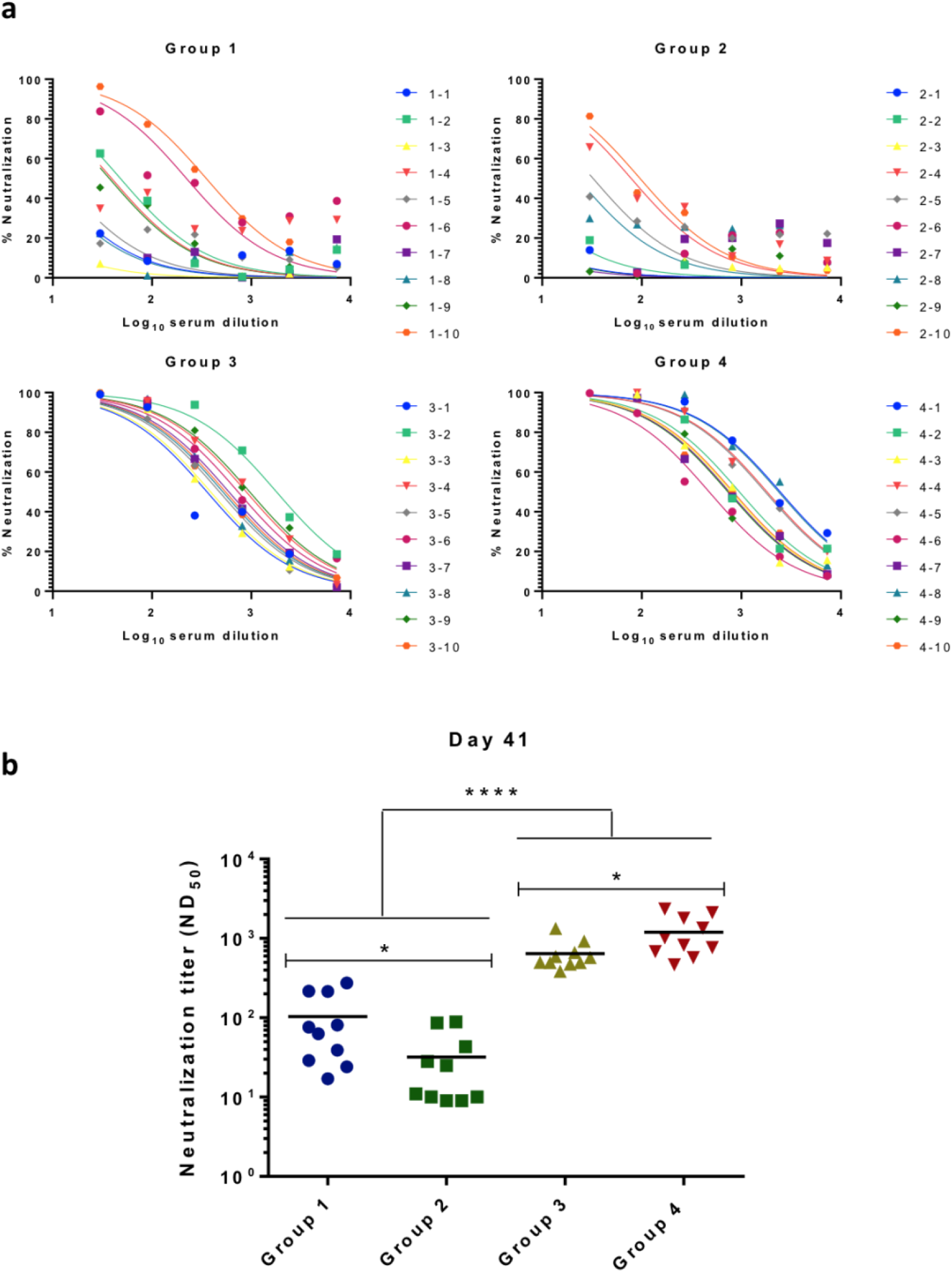
Neutralization study of mice sera on SARS-CoV-2 pseudo-typed virus. (a) Curves showing the percentage of neutralization of individual sera of mice from all groups on viral infection by SARS-CoV-2 pseudo-typed virus. This pseudo-typed viruses encodes luciferase as a reporter of cell infection and was preincubated with three-time serial dilutions of sera. Luciferase expression was compared to the one of non-neutralized virus. (b) Comparison of the neutralization titer at 50% of the maximal effect (ND50) from all individual mice with the mean represented by a line. Statistical analysis demonstrates the superiority of neutralization of sera from mice immunized with RBD-decorated ADDomer particles.

## Discussion

In this study, we designed a novel and highly versatile vaccine platform amenable to large antigens by adapting the SpyTag/SpyCatcher technology to ADDomer, an adenovirus-inspired virus-like particle (VLP)^18,19^. This technology has already been described for designing VLP-derived vaccine, however in these approaches SC was the moiety grafted to the VLPs while the antigens were fused to the Spy Tag^20,24^. In our novel vaccine technology, we used the opposite configuration with SpyTag insertion in an internal loop of ADDomer, which minimized the risk of steric hindrance and functional impact on the spontaneous oligomerization of ADDomers. Insertion of ST in a surface-exposed loop of ADDomer did not impair the spontaneous assembly of twelve pentameric bricks (Figure 1d), thus creating a universal generic platform for multivalent antigen display. Experiments of saturating cross-linking with SC alone suggested that a full decoration (*i*.*e*. 60 copies) of the particle can be achieved (Figure 1b). Despite the flexibility of the loop in which ST was inserted, cryo-EM reconstruction of the ADD-ST/SC complex clearly showed an extra density corresponding to the SC interacting with ST fused to the ADDomer. The data thus strongly suggested that the ADDomer could be used to display a multimerized array of antigens, a property that is crucial for the elicitation of strong antibody responses^25,26,27^. Interestingly, the characteristic 2, 3 and 5-fold symmetry axis of a dodecahedron are easily visible in the 3D structure (Figure 2c), this spatial configuration showing that the SCs are distributed at different distances from each other according to the symmetry axis. Such distribution of the antigen could potentially further favor the immunogenicity of the VLP by offering different patterns of presentation of the antigen at the particle surface.

The COVID pandemic led us to first evaluate our newly designed platform as an anti-SARS-CoV-2 vaccine, using the RBD as the antigen to be displayed through fusion to the SC N-terminus. It is worth noting that in the 3-fold axis of ADDomer, the SC adopts a pseudo-trimeric arrangement, which on the RBD decorated VLP closely mimics the tridimensional configuration of this domain in the SARS-CoV-2 spike glycoprotein (Figure 2d). The distance between SC in such configuration is 4,7 nm, which is in the same range as that between RBDs in the trimeric Spike protein (4,1 nm). Expressing soluble RBD fused to SC in insect cells resulted in N-glycosylation as expected^28^ and did not affect the structure of SC, which retained its ability to cross-link to the ADD-ST vaccine platform in a dose dependent manner (Figure 3c). The proper folding of the glycosylated RBD exposed at the surface of the particles and its ability to bind ACE2 were assessed in a functional viral entry assay *via* competition with SARS-CoV-2 Spike pseudo-typed virions (Figure 4c). A surface plasmon resonance (SPR) experiment confirmed the high affinity interaction of ADD-RBD to ACE2, the highly stable interaction observed suggesting an avidity effect of the multivalent particle exposing multiple copies of RBD (Figure 4a).

The ability of the RBD-decorated ADDomer to elicit an anti-SARS-CoV-2 humoral response was then studied in mice. As expected, we confirmed that displaying the RBD in a multimerized manner led to a strong improvement in immunogenicity (groups 3 and 4 *versus* groups 1 and 2). Indeed, this was clearly apparent from the first injection where the groups with decorated particles showed a significant anti-RBD response at Day 13 as measured in ELISA (Figure 5c), whereas no response was observed for the other groups. After the second immunization, although a response was observed in the non-decorated groups, the titers attained in the decorated groups were about 10 fold greater (Figure 6a). Notably, the anti-RBD response in the latter groups endured over time with similar titers at 2 and 4 weeks following the second immunization (Figure 6b). The anti-SARS-CoV-2 RBD sera were further assessed for their capacity to neutralize SARS-CoV-2 in pseudo-typed virus-based assay (Figure 7). As seen for binding responses, a neutralizing activity was observed in the RBD-decorated groups after the first immunization. Overall, a good correlation between ELISA and neutralizing titers was observed. Strong neutralizing titers were attained in all mice immunized with ADD-RBD after the boost immunization, showing the ability of our vaccine platform to elicit functional antibody responses with a potential protective activity. The low titers obtained in mice immunized with RBD and naked ADDomer (no decoration) again demonstrated that the high immunogenicity achieved with the decorated VLPs originated from the antigen multimeric display. The data are in adequacy with results obtained with similar vaccine platforms using multimeric display of the SARS-CoV-2 RBD on scaffolds such as the 24mer-ferritin, the 60-mer dodecahedral thermophilic aldolase ^24,29,30^. In contrast to these latter vaccine platforms, our particles originate from a human virus, Adenovirus serotype 3 (HAd3), which is known to have a relatively high seroprevalence^31,32^. A potentially detrimental impact of preexisting immunity on the immunogenicity of adenovirus-based viral vectors has been described^33^, which thus led us to explore whether such effect may also be encountered with our adenovirus-derived VLP platform. Interestingly, in contrast to adenovirus-viral vectors, our adenovirus-inspired VLP platform appears to benefit from pre-immunity. Indeed, mice with immunity to ADDomer at the time of the first immunization were found to respond with greater antibody titers against RBD than naive mice (group 4 *versus* group 3, Figure 5c). This beneficial effect may be in particular explained by the formation of immune complexes leading to a better antigen presentation increasing immunogenicity^34^. This property persisted after the second immunization although the differences between groups 3 and 4 were less marked, probably due to an anti-ADDomer response generated in group 3 during the first injection.

In conclusion, we designed a new adenovirus-inspired vaccine platform to achieve high immunogenicity against the displayed antigen. This platform efficacy is not dependent on the immunological status against adenovirus which allows to consider its use in a large coverage. We exemplified in the present work the ability of this technology to generate SARS-CoV-2 potent neutralizing antibodies against the Wuhan strain, but its ease of use allows to envision a rapid application to different variants of concern (VOCs). Moreover, the versatility of the vaccine platform paved the way to generate mosaic particles displaying RBDs from different betacoronaviruses^35^or from different SARS-CoV-2 variants at once. More generally, this novel platform constitutes a precious tool in the preparedness to the probable emerging pathogens.

## Material and Methods

### Baculovirus production

The baculovirus expression system was used for both the production of ADDomer-SpyTag (ADD-ST) and for the RBD fused to SpyCatcher (RBD-SC). Synthetic DNA (Genscript) was cloned in the pACEBac1 using the restriction sites BamHI and HindIII. For RBD-SC, the SARS-CoV-2 spike sequence (320-554) was cloned upstream of the Spy Catcher and a hexa His-Tag was added in C-terminus of SC. This fusion protein was secreted using the melittin signal peptide present in the vector. Recombinant baculoviruses were made by transposition with an in house bacmid expressing Yellow Fluorescent Protein, as previously described^18^. Baculovirus were amplified on Sf21 cells at low multiplicity of infection (MOI) and after two amplification cycles were used to infect insect cells for 64-72h at high MOI. For ADD-ST production, the infected cells were pelleted and recovered whereas for RBD-SC cells were discarded and the supernatant was saved.

### Protein Purification

#### ADDomer and ADD-ST purification

The ADDomer and ADD-ST was purified according to classical protocol^36^. Briefly, after lysis of the insect cell pellet by 3 cycles of freeze-thaw in the presence of Complete protease inhibitor cocktail (Roche), and removal of debris the lysate was loaded onto a 20-40% sucrose density gradient. The gradient was centrifuged for 18h at 4°C on a SW41 Rotor in a Beckman XPN-80 ultracentrifuge. The dense collected fractions at the bottom of the tubes were dialyzed against Hepes 10 mM pH 7.4, NaCl 150 mM and then loaded onto a Macroprep Q cartridge (Bio-rad). After elution by a 150 to 600 mM linear NaCl gradient in Hepes 10mM pH 7.4, ADDomer-containing fractions were checked by SDS-PAGE and concentrated on Amicon (MWCO: 100kDa) with buffer exchange to Hepes 10 mM pH 7.4, NaCl 150 mM.

#### Spy Catcher (SC) purification

After lysis of the insect cell pellet as described above in the presence of Complete EDTA-free protease inhibitor cocktail, the clarified lysate was diluted 5-time in wash buffer (Hepes 10mM pH7.4, NaCl 150mM, imidazole 10mM and loaded onto a His Gravitrap column (Cytiva) by gravity, with two passages of the lysate onto the column. The column was washed with the same buffer, then eluted by 200 mM Imidazole. The fractions were analyzed by SDS-PAGE, pooled and Imidazole was withdrawn by buffer exchange using Amicon ultrafiltration devices (MWCO: 4 kDa).

#### RBD-SC purification

The insect cell supernatant was centrifuged after thawing for 15 min at 7,500 g and loaded onto a Hepes 10 mM pH7.4 pre-equilibrated Heparin column (Cytiva) of 5 ml for 500 ml of supernatant. The column was washed with Hepes 10 mM pH 7.4 for 25 mL then eluted for 10 mL with 0 to 500 mM linear NaCl gradient in Hepes 10 mM pH 7.4. The eluate was supplemented with 30 mM Imidazole-HCl and incubated with Ni-NTA beads (Qiagen), 2 ml of beads for 500 ml culture, for at least 1h at 4°C under gentle agitation. The beads were then poured into an empty column and the protein was eluted by 2 column volumes of 250 mM Imidazole in Hepes 10 mM pH 7.4, NaCl 150 mM. It was then submitted to buffer exchange using Amicon device (MWCO 30 kDa).

#### RBD purification

The following reagent was produced under HHSN272201400008C and obtained through BEI Resources, NIAID, NIH: Vector pCAGGS Containing the SARS-Related Coronavirus 2, Wuhan-Hu-1 Spike Glycoprotein Receptor Binding Domain (RBD), NR-52309.Vector NR-52309 from BEI Resources was used for mammalian expression of the RBD alone used for ELISA. EXPI293 cells grown in EXPI293 expression medium were transiently transfected with the vector according to the manufacturer’s protocol (Thermo Fisher Scientific). Five days after transfection, the medium was recovered and filtered through a 0.45 mm filter. Two-step protein purification on Aktä Xpress, with a HisTrap HP column (GE Healthcare) and a Superdex 75 column (GE Healthcare) was performed using 20 mM Tris pH 7.5 and 150 mM NaCl buffer. For the HisTrap, a wash step in 75 mM imidazole was performed and RBD was eluted in buffer supplemented with 500 mM imidazole before loading onto the gel filtration column run in equilibration buffer.

#### N-glycosidase treatment

RBD-SC was incubated for 1H at 37°C with N-glycosidase (kindly provided by Dr Nicole Thielens) at 1/100 ratio then run on gel next to the same amount of untreated RBD-SC.

### Complex formation

#### ADD-SpyTag + SpyCatcher (ADD-ST/SC) for Cryo-EM

The purified ADD-ST was mixed with an excess of purified SpyCatcher (ratio 1:4). The protein mix was incubated overnight at 25°C under shaking in Thermomixer (300 rpm). A purification step on a sucrose gradient was performed to remove the SpyCatcher in excess and to recover only the fully-decorated ADD-ST/SC at the bottom of the gradient. Buffer exchange was done in10 mM Hepes, 150mM NaCl and the sample was concentrated to 1.5 mg/ml.

#### ADD-SpyTag + SpyCatcher-RBD (ADD-RBD) for characterization and mice immunization experiments

Covalent complex formation was obtained by incubation of purified ADD-ST with purified RBD protein fused to SC (RBD-SC). Incubation was performed at 25°C under agitation on a Thermomixer at 300 rpm. RBD-SC ratio per ADD-ST was varied according to the experiment as indicated in the text. For immunization experiments, a ratio of 40 copies of RBD-SC per ADD-ST was chosen. This ratio was calculated on SDS-PAGE using ImageLab software (Bio-Rad). Integrity of the ADD-RBD was checked by negative staining electron microscopy.

### In vivo experiments and vaccination

Vaccination experiments have been performed according to ethical guidelines, under a protocol approved by the Grenoble Ethical Committee for Animal Experimentation and the French Ministry of Higher Education and Researcher (reference number: APAFIS#27765-2020102114206782 v2). 5-week-old female Balb/c mice were purchased from Janvier (Le Genestet St. Isle, France). For all mice groups, vaccines were in PBS and adjuvantized with one volume of ADDavax (Invivogen). All the mice received 2 vaccine doses at 2 weeks of interval (Day 0 and Day 14) starting at 8-weeks of age. Each mice group received subcutaneously the corresponding vaccine as indicated within the paper in 100µl final volume, in the right flank. Pre-immunized group received 2 weeks before the first dose of vaccine (day -15) the ADDomer vector alone (5µg in 100 µl final volume, in the right flank). The day before each vaccination, and 2 and 4 weeks after the last vaccination, 100µl of blood was withdrawn for serologic tests. For retro-orbital blood sampling, mice were anesthetized with 4% Isoflurane.

### Electron microscopy

#### Negative staining

3.5 μL of samples were adsorbed on the clean side of a carbon film previously evaporated on mica and then stained using 2% (w/v) Sodium Silico Tungstate pH 7.4 for 30 s. The sample/carbon ensemble was then transferred to a grid and air-dried. Images were acquired under low dose conditions (<30 e−/Å2) on a F20 electron microscope operated at 120 kV using a CETA camera.

#### Cryo-EM

Quantifoil grids (300 mesh, R 1.2/1.3) were negatively glow-discharged at 30 mA for 45 s. 3.5 µl of the sample were applied onto the grid, and excess solution was blotted away with a Vitrobot Mark IV (FEI) (blot time: 6 s, blot force: 0, 100% humidity, 20 °C), before plunge-freezing in liquid ethane. The grid was transferred onto a 200 kV Thermo Fisher Glacios microscope equipped with a K2 summit direct electron detector for data collection. Automated data collection was performed with SerialEM, acquiring one image per hole, in counting mode. Micrographs were recorded at a nominal ×36,000 magnification giving a pixel size of 1.145 Å (calibrated using a β-galactosidase sample) with a defocus ranging from −0.6 to −2.35 µm. In total, 1038 movies with 40 frames per movie were collected with a total exposure of 40 e−/Å^2^.

#### Image processing and cryo-EM structure refinement

Movie drift correction was performed with Motioncor2 ^37^ using frames from 2 to 40. CTF determination was performed with Relion 3.1.2 (Scheres 2012). 939 movies out of 1038 were kept at this stage. Particles selection have been done using the Laplacian filter with a diameter between 30 and 40 nm. A total of 77 323 particles have been automatically selected, boxed into 480×480 pixel^2^ boxes and submitted to 2D classification. After extensive selection and generation of an initial model imposing I1 symmetry, 3D refinement followed by CTF Refine and post-processing generated a final reconstruction including 10163 particles with a resolution of 2.76 Å (Fourier Shell = 0.143) (applied B-factor -82)

### Surface plasmon resonance

Surface plasmon resonance experiment was performed on a T200 instrument. Anti human Fc polyclonal antiboby (Jackson Immunoresearch, 109-005-008) diluted at 25 µg/ml in 10 mM sodium acetate pH 5 was immobilized on CM5 sensor chips using the amine coupling chemistry according to the manufacturer’s instructions (Cytiva) to get an immobilization level of 14 000 RU. ACE-2 –Fc (GenScript Z033484) was diluted at 1.2µg/ml in HBS P+ (Cytiva) to get an capture level of 100RU. For interaction measurements, ADD-RBD (ranging from 1 nM to 11nM in HBSP+) was injected over captured ACE-2 Fc in HBS P+ buffer at 30 µl/min. Anti human Fc polyclonal antiboby flow cell was used for correction of the binding response. Regeneration of the surfaces was achieved by 10mM Glycine pH2. Binding curves were analyzed using BIAEvaluation software (GE Healthcare) and data was fit to a 1:1 Langmuir with drifting baseline interaction model.

### Neutralization assays and pseutdo-typed SARS-CoV-2 virion production

#### Pseudovirus production and titration

Neutralization assays were performed using lentiviral pseudotypes harbouring the SARS-CoV-2 spike and encoding luciferase. Briefly, gag/pol and luciferase plasmids were co-transfected with a SARS-CoV-2 spike plasmid with a C-term deletion of 18aa at 1/0,4/1 ratio on adherent HEK293T cells. Supernatants containing the produced pseudoviruses were harvested 72H after transfection, centrifuged, filtered through 0.45µm and concentrated 50 times on Amicon Ultra (MWCO 100KDa), aliquoted and stored at -80°C. Before use, supernatants were tittered using HeLa ACE-2 cells to determine the appropriate dilution of pseudovirus necessary to obtain about 150 000 RLU per well in a 48 well-plate.

#### Neutralization assay (mAb)

Serial 3-fold dilutions starting from 1/10 dilution (serum), or 6µg/mL (known bNAbs) were let in contact with the pseudoviruses for 1h at 37°C in 96-w white plates (Greiner #675083), before addition of HeLa ACE2 cells. Plates were incubated 24h at 37°C, protected from evaporation, then cells were fed with 60 µL of DMEM (Gibco 11966-025) supplemented with 10% FBS (VWR 97068-086), and incubated for another 24h. Medium in each well was aspirated and replaced by 45 µL of 1X cell lysis buffer (Ozbioscience # LUC1000) for a 60 min-incubation under agitation. Thirty µL of luciferin substrate was then added and Relative Light Unit (RLU) was measured instantly by a luminometer.

#### Statistical analyses

For ELISA and neutralization assays, IC50 were deduced from crude data after normalization using GraphPad Prism “log(inhibitor) vs normalized response” function. Comparison of IC50 values were performed applying Mann-Whitney tests on the data sets, using GraphPad Prism 6 software.

## Acknowledgments

We are grateful to the ‘CNRS Prematuration program’, to the ‘ANR-Flash COVID’ and to ‘Région Auvergne Rhône-Alpes, AuRA’ for the financial support to this work. This project has received funding from the European Research Council (ERC) under the European Union’s Horizon 2020 research and innovation program (grant agreement No 682286). D.H is supported by GEFLUC Dauphiné-Savoie, Ligue contre le Cancer Comité Isère, Université Grenoble Alpes IDEX Initiatives de Recherche Stratégiques and Fondation du Souffle-Fonds de recherche en santé respiratoire (FdS-FRSR). We thank Aymeric Peuch for help with the usage of the EM computing cluster and Emmanuelle Neumann for the training of CC in negative staining and Leandro Estrozi for help with image analysis. This work used the platforms of the Grenoble Instruct-ERIC centre (ISBG ; UAR 3518 CNRS-CEA-UGA-EMBL) within the Grenoble Partnership for Structural Biology (PSB), supported by FRISBI (ANR-10-INBS-05-02) and GRAL, financed within the University Grenoble Alpes graduate school (Ecoles Universitaires de Recherche) CBH-EUR-GS (ANR-17-EURE-0003). The electron microscope facility is supported by the Auvergne-Rhône-Alpes Region, the Fondation Recherche Medicale (FRM), the fonds FEDER and the GIS-Infrastructures en Biologie Sante et Agronomie (IBISA). IBS acknowledges integration into the Interdisciplinary Research Institute of Grenoble (IRIG, CEA). V.D is an internship of Master de Biologie, École Normale Supérieure de Lyon, Université Claude Bernard Lyon I, Université de Lyon, 69342 Lyon Cedex 07, France.

## Author Contributions

P.F, M.C.D, PP conceived the experiments. Vector production and purification were performed by C.C, V.D and S.G. Structural analysis was performed by G.S, D.F and C.C. Animal experiments were performed by D.H and S.B. Molecular interaction studies was performed by E.G, C.M and E.V.S. Immunological characterization and virus neutralization was performed by A.A, I.B. and M.B With input from all authors, P.F and P.P wrote the manuscript.

## Competing interests

We declare no competing interests

## Notes

### Competing Interest Statement

The authors have declared no competing interest.

