## Supplementary Figures for "Elicitation of potent SARS-CoV-2 neutralizing antibody responses through immunization using a versatile adenovirus-inspired multimerization platform"

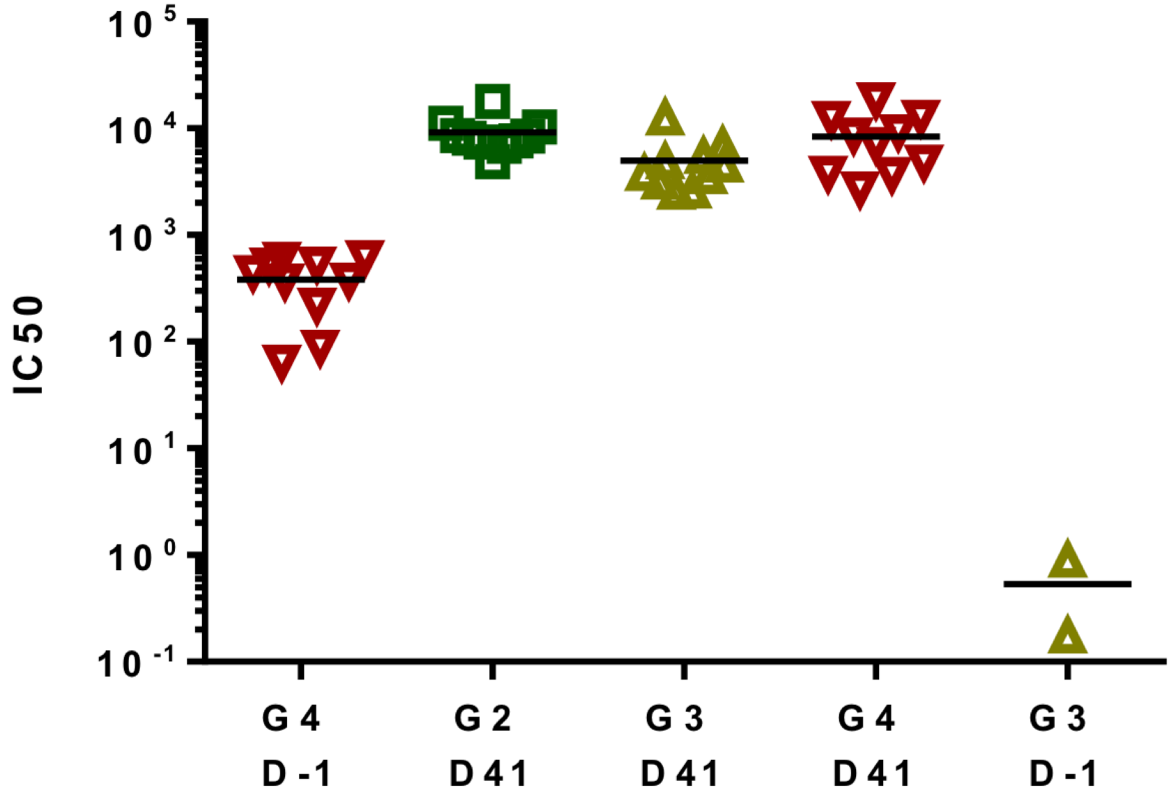


**Supplementary Figure 1: Anti-ADDomer response studied by ELISA.**
Non-decorated ADDomer were coated on the well and incubated with sera from different groups of mice before the first injection (D-1) or at the end of the experiment (D41). Note that all the mice preimmunized with ADDomer (G4, D-1) were positive before injection of ADD-RBD while naïve mice (G3, D-1) were negative as expected.

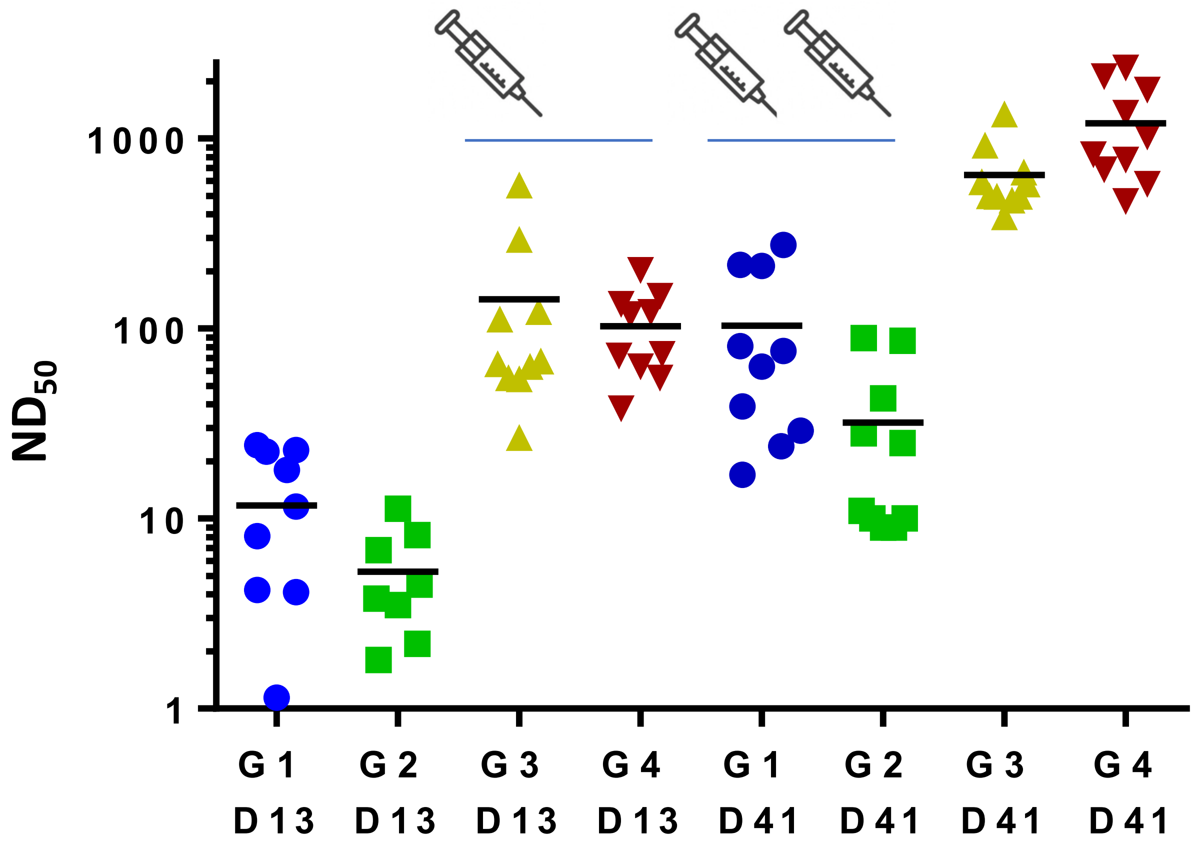


**Supplementary Figure 2: Sera neutralization titer after one and two injections for all groups of mice.** Note that RBD displayed on ADDomer enabled to get higher neutralization titer at the first injection (G3 and G4, D13, depicted with 1 syringe) than RBD not displayed on the platform after two injections (G1 and G2 at D41, depicted by 2 syringes)
